# RNA-Protein Interaction Prediction without High-Throughput Data: An Overview and Benchmark of *in silico* Tools

**DOI:** 10.1101/2024.06.24.600368

**Authors:** Sarah Krautwurst, Kevin Lamkiewicz

## Abstract

RNA-protein interactions (RPIs) are crucial for accurately operating various processes in and between organisms across kingdoms of life. Mutual detection of RPI partner molecules depends on distinct sequential, structural, or thermodynamic features, which can be determined via experimental and bioinformatic methods. Still, the underlying molecular mechanisms of many RPIs are poorly understood. It is further hypothesized that many RPIs are not even described yet. Computational RPI prediction is continuously challenged by the lack of data and detailed research of very specific examples. With the discovery of novel RPI complexes in all kingdoms of life, adaptations of existing RPI prediction methods are necessary. Continuously improving computational RPI prediction is key in advancing the understanding of RPIs in detail and supplementing experimental RPI determination. The growing amount of data covering more species and detailed mechanisms support the accuracy of prediction tools, which in turn support specific experimental research on RPIs. Here, we give an overview of RPI prediction tools that do not use high-throughput data as the user’s input. We review the tools according to their input, usability, and output. We then apply the tools to known RPI examples across different kingdoms of life. Our comparison shows that the investigated prediction tools do not favor a certain species and equip the user with results varying in degree of information, from an overall RPI score to detailed interacting residues. Furthermore, we provide a guide tree to assist users which RPI prediction tool is appropriate for their available input data and desired output.

## Introduction

Interactions between RNAs and proteins (RPIs) are essential for multiple molecular processes in biological entities. Key players for RPIs are RNA-binding proteins (RBPs), which are involved in gene expression, RNA processing, modification, and degradation of RNA [1–3]. Interaction of mRNAs with RBPs can initiate or regulate protein synthesis [4–6]. The function of microRNAs (miRNAs) (for the regulation of gene expression) depends on RBPs [7, 8], same as for long non-coding RNAs (lncRNAs) [9, 10]. Due to the vast variety of processes in which RPIs are involved, their functionalities are also linked to diseases [11–13].

Most knowledge on RBPs originates from eukaryotic systems, especially the human organism [14], for which around 1,500 RBPs are annotated [15]. However, data for bacteria and viruses is sparse, as for a typical bacterium, around 180 RBPs are known [16]. For viruses, most RPI research focuses on host-virus rather than intra-viral RPIs, which started to set off only a decade ago [17].

RPIs are realized dynamically with a one-sided or mutual conformational change of the RNA and protein partner [18–20]. Additionally to this conformational change, many residues may not be part of the interaction directly but are still crucial for binding site flexibility and correctly positioning functional residues in RPIs [21, 22]. RBPs bind RNA molecules either specifically (e.g. based on RNA modifications [23], sequential or structural motifs [24, 25]) or non-specifically (e.g. dsRNA or ssRNA in general [24, 26, 27]). For the protein, aromatic and positively charged amino acids are often involved in contacting the RNA partner, especially since they can form specific and strong interactions like salt bridges and *π*-stacking with the nucleobases [24, 28]. However, the backbone of the RNA is associated more often with the protein than the bases in RPIs [29]. Solvent accessibility and structural positions of the amino acids are decisive for interaction as well [21]. Usually, RNA binding domains (RBDs) are the main interaction area. Single amino acids outside of the RBD can take further action toward contacting the RNA [1, 28].

Experimental detection (*in vitro* and *in vivo*) of RPIs [30, 31] can focus on an RNA molecule of interest to characterise potential proteins binding to it (e.g. RAT/TRAP, RNA affinity in tandem / tagged RNA affinity purification [32, 33]) or on a known RBP to identify interacting RNAs (RIP-Chip, RNA immunoprecipitation chip [34]; CLIP, cross-linking immunoprecipitation [35]). *In vitro* methods can result in RPIs that are not physiologically relevant [1], which can be avoided by using *in vivo* approaches. Non-crosslinking *in vivo* methods like RIP-Chip [34], which combines immunoprecipitation with RT-PCR and microarrays, can come along with noise issues, co-immunoprecipitation of unwanted additional RBPs, false positives from re-associated RBP and RNAs after cell lysis [36, 37], or no possibility to identify the binding site specifically because of mild conditions for preserving the non-covalent RPIs [38]. Instead of microarrays, RIP-Seq combines RNA immunoprecipitation with high-throughput sequencing [39]. Alternatives are CLIP-based (crosslinking and immunoprecipitation) methods (e.g. HITS-CLIP, iCLIP, Par-CLIP), which solve some of the former issues and carry other disadvantages. For example, HITS-CLIP (high-throughput sequencing CLIP) enables large-scale RPI detection [40], but the UV radiation from the UV-crosslinking step prior to immunoprecipitation may lead to mutagenesis [41], or specificity issues during crosslinking (biased for ssRNA, pyrimidines, certain amino acids) [38, 42]. Par-CLIP (Photoactivatable-Ribonucleoside-Enhanced CLIP) [43] and iCLIP (individual nucleotide resolution CLIP) [44] both provide specific resolutions of interaction-involved nucleotides and binding sites but require long technical procedures (Par-CLIP [45]) and demanding set-ups (iCLIP [46]), involving numerous reaction steps. In general, *in vivo* techniques grant biologically more specific (e.g. regarding tissue/cell lines) interaction data, which is beneficial for specific training of computational RPI prediction algorithms. The potentially high false negative [47, 48] and false positive [49, 50] rate of these experiments however could lead to biases in training of such algorithms.

Although the above problems of experimental RPI detection have mostly been tackled with modern and up-to-date experimental workflows, they can still be expensive, time-consuming, or require challenging set-ups [30]. Furthermore, meeting the exact conditions for the RPIs to happen and be detectable *in vivo* (e.g. tissue-specificity, cell-cycle specificity, time-dependency, robustness of interaction) can prove difficult [51]. Therefore, experimental approaches can be supplemented with bioinformatics methods. Algorithms help analyze existing experimental data sets in more detail [52, 53], investigate available sequences and structures for motifs and properties [54], predict binding sites of RNA-protein partners based on similar known interactions [55], or classify whether an RPI is probable [56]. Input for the algorithms is mostly the sequence or PDB (Protein Data Bank) [57] 3D structure of either or both RPI partners. The output ranges widely between tools, e.g., getting an overall interaction score for an RNA and protein pairing [58, 59], sequence motifs contributing to an RPI [52, 60], a potential binding site area highlighted in a protein structure [61–63], or specific interacting residues between an RNA-protein complex [64]. Many RPI prediction tools are available, varying in aspects like background data, feature selection, machine-learning algorithm, or output extent. In recent years, several tools and workflows have been proposed that utilize experimental data from, e.g., CLIP-Seq experiments. While these resources are valuable and often used to determine specific binding motifs or residues of a protein of interest, there are scientific questions or use cases where such data is unavailable. For an overview of tools available that use high-throughput sequencing (HTS) data, we refer to [65–67].

Here, we present a comparative analysis of RPI prediction tools that do not need experimental HTS data as input. First, we provide a comprehensive overview of available tools grouped by their input requirements. With many RPI prediction tools being developed in recent years, users might lose the overview of accessible tools fitting their needs. To assist potential users, we propose a guide tree covering available RPI prediction algorithms to identify tools of interest for different applications and use cases.

Furthermore, many RPI prediction tools are often benchmarked or usable only with HTS data sets. However, since such data is rarely available for non-model organisms or users might only be interested in a specific RNA-protein complex, we focus on tools that allow such a single-RNA-protein-complex as input. We apply the 30 collected available algorithms on four selected RPI examples across different kingdoms to assess a potential bias towards specific taxonomic clades. Thus, we focus on (i) the human protein LARP7 binding to the 7SK snRNA; (ii) the MS2 phage coat protein interacting with an RNA hairpin in the phage’s genome; (iii) the Ebola virus VP30 protein binding to the viral RNA leader region; and (iv) the bacterial toxin-antitoxin system ToxIN. Using this small subset of known RPI examples and their respective “ground truth” of interactions from literature, our evaluation provides a more detailed insight into the capabilities and applicabilities of the RPI prediction algorithms.

## Results

*De novo* RPI prediction tools need sequence or structure input of the potentially interacting RNA or protein molecules. Predicted results vary in the degree of information, e.g., some tools only report interaction scores, whereas others report motifs, energies, binding sites, or interacting residues. To assist users in deciding what tool to use for their RPI prediction analysis, we compiled an extensive overview of available RPI prediction tools at GitHub. We further summarized this overview in a guide tree shown in Fig. 1. The tree covers currently (at the time of writing) accessible RPI prediction tools, categorized into necessary input data, computed output, and whether a web server or stand-alone version is available. This overview also contains prediction tools relying on experimental HTS data (see Fig. 1, input data: ”experimental dataset”), which were not part of our evaluation. In the following, we review the collected 30 *de novo* RPI prediction tools (see Fig. 1, input data: ”RNA”, ”protein”, ”RNA and protein”) and subsequently present the evaluation results for the available algorithms.

**Figure 1:**
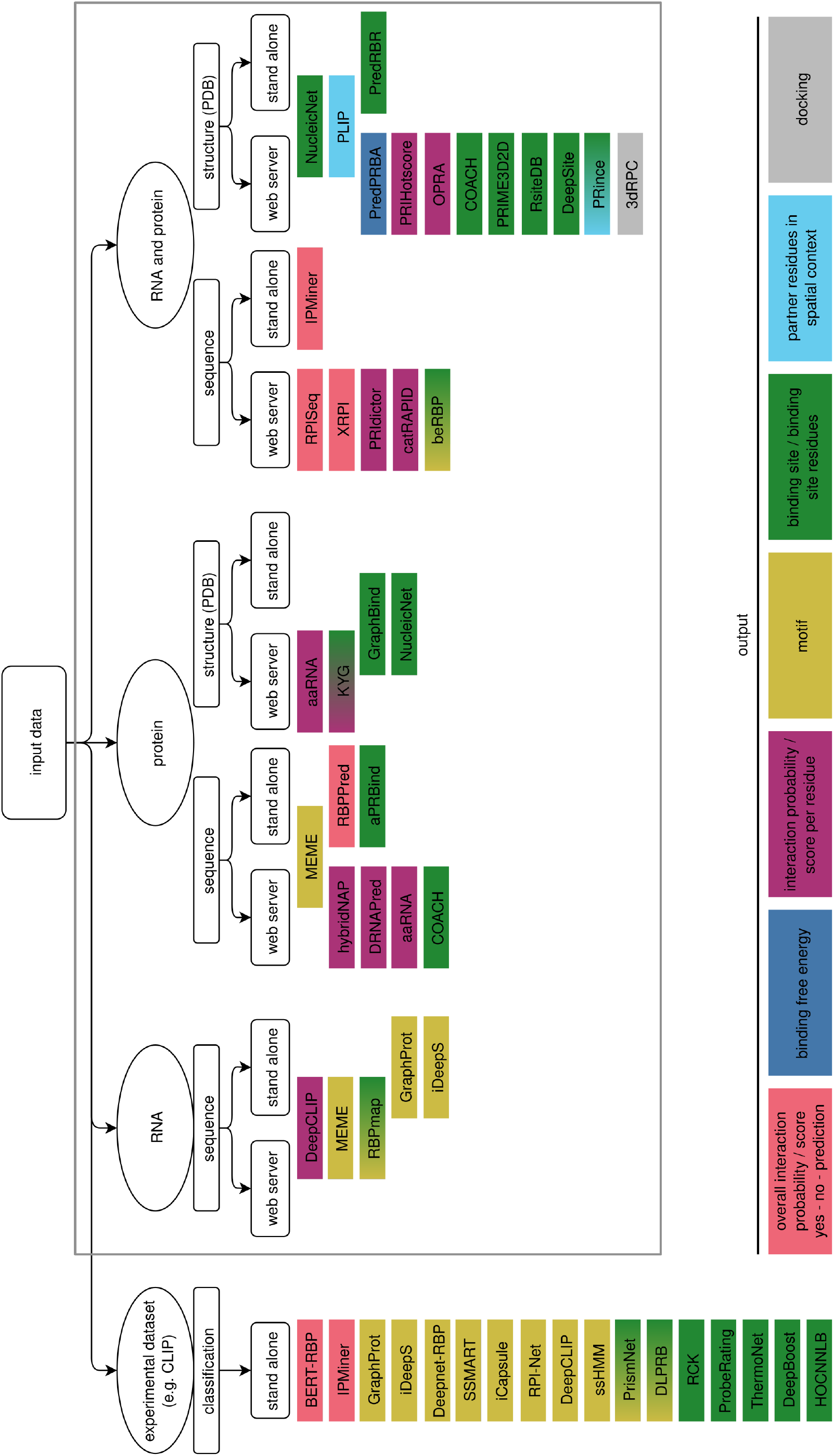
Guide tree for available RPI prediction tools given specific input data. We provide a (non-exhaustive) list of tools that are still maintained or are at least accessible at the time of writing. Additionally to the input, tools are categorized based on web server or stand-alone versions. The respective output type is color-indicated. A comprehensive tabular overview is also given at GitHub. All tools within the gray-framed box are part of our evaluation in this study. Figure created with InkScape [68].

### Overview of *de novo* RPI prediction tools

#### RNA sequence input

In general, tools using the RNA sequence alone, such as MEME [69], GraphProt [70], and iDeepS [71], report putative sequence motifs of the RNA involved in the interaction. It should be noted that these three tools also have further input options (protein sequence for MEME, HTS datasets for Graph-Prot and iDeepS). MEME is a motif discovery tool, requiring any kind of biological sequences as input, and calculating (*de novo*) sequence motifs therein, which are displayed and listed for the user. Although not specifically targeted for RPIs, MEME can still serve as an initial step in analyzing potential RNA and protein partners [69]. While both GraphProt and iDeepS work with RNA sequence input, usage still depends on an own HTS dataset to train or the availability of a matching pre-trained model (24 human RBPs for GraphProt, 31 for iDeepS), which was not the case for our chosen biological examples. Therefore, we excluded these two tools from our evaluation. DeepCLIP [72] is a deep neural network trained on CLIP datasets, however, it expects RNA sequence(s) as input, if the user seeks to work with one of the pre-trained models. DeepCLIP calculates RBP binding probabilities for the positions of the given input [72]. RBPmap [60] requires RNA sequence(s) as input as well. Additionally, RBPmap is able to search for RBP-binding motifs in the query RNA(s), which the user has to provide. The motifs are restricted to RBPs in human and two model organisms (*Mus musculus, Drosophila melanogaster*).

#### Protein sequence input

If only protein data is available, users can refer to hybridNAP [73], DRNAPred [74], or aaRNA [75]. These tools expect the sequence of the protein and report individual interaction probabilities or scores per amino acid. hybridNAP considers potential binding residues by their relevant sequential properties, relative solvent accessibility in the structure, and their evolutionary conservation, which are assessed based on their test data [73]. These features are covered by DRNAPred as well, in addition to putative intrinsic disorder and secondary structure [74]. The algorithm of aaRNA furthermore includes homology [75]. Similarly, aPRBind [76] annotates residues potentially involved in an interaction using features from protein structure models additionally to the protein sequence, thus trying to incorporate dynamic properties relevant for the potential interaction [76].

Aside from an RNA sequence, MEME can also process protein sequences and reports motifs of interest on the protein. RBPPred [77] allows for the rapid scanning of many proteins by utilizing an SVM classifier to predict whether a protein can bind to an RNA by considering evolutionary information and physicochemical properties of the primary sequence. For each input sequence, the algorithm predicts whether it can bind RNA or not [77].

#### Protein structure input

Given the structure of a protein (most commonly in PDB-like format), GraphBind [78], NucleicNet [63], and KYG [79] are available to predict the binding sites for RNA partners in the protein structure. Users can choose GraphBind to calculate interactions specific for different kinds of ligands. For the prediction, graphs are constructed to reflect the structure context and important features, which are then fed into hierarchical graph neural networks (HGNNs) [78]. NucleicNet predicts and visualizes whether RNA can be bound across the grid of the protein surface given the physicochemical environment, specifically the interaction modes for the different parts of RNA molecules [63]. Furthermore, general binding potential of RNA sequences can be evaluated using logo diagrams. The algorithm is based on the FEATURE vector framework [80], with feature vectors encoding the physicochemical properties. KYG calculates interface residue propensities for each amino acid and residue pairing preferences between the protein and RNA, using data of representative RNA-protein complexes from the PDB [79].

Furthermore, aaRNA can work with a protein structure as well, which results in a structural visualization additionally to the per-residue-probability prediction [75].

#### RNA and protein sequence input

The highest prediction accuracy is possible when RNA and protein information is available. On sequence level, tools such as XRPI [59], RPISeq [58], and IPMiner [81] provide an overall interaction probability, whereas PRIdictor [55] and catRAPID [82] report individual probabilities for residues potentially involved in the interaction. XRPI is a machine-learning method, calculating the interaction probability via a gradient boosting classifier (XGBoost) based on features, such as smallest structural unit and amino acid interaction propensities, from RNA-protein structures in the PDB [59]. The predicted interaction score ranges from 0 to 1. Similarly, RPIseq [58] provides two such scores for a potential RNA and protein sequence pair, calculated by classical machine-learning approaches as well. These two respective classifiers (Support Vector Machine (SVM) and Random Forest (RF)) are trained on non-redundant datasets from the Protein-RNA Interface Database (PRIDB) [83]. The deep learning tool IPMiner extracts features by a stacked autoencoder and uses those for prediction via stacked ensembling of three random forest classifiers [81].

PRIdictor calculates the mutual binding site residues using global and local features of the sequences, as well as partner features, encoded in feature vectors. The predictions consider hydrogen bonds, water bridges, and hydrophobic interactions as potential RPIs [55]. Based on the contributions of hydrogen bonding, van der Waals contacts, and predicted secondary structure of the RNA and protein domains, catRAPID calculates interaction propensities for RNA and protein [82]. With its training data, the algorithm of catRAPID computes and visualizes the pairwise interaction scores for all residues of a given RNA-protein pair and the corresponding discriminative power in a heatmap.

The model of beRBP [84] predicts RBP-binding site motifs in given input RNA sequence(s). The user additionally has to provide or choose a pre-trained position weight matrix (PWM) model for a RBP, or a RBP sequence, in which case beRBP tries to predict a corresponding PWM based on similarity of potential RBDs. The ‘General model’ is trained with a Random Forest approach based on a matrix considering four features of a putative binding site: the match of a motif, sequence environment, spatial accessibility and evolutionary conservation [84].

#### RNA and protein structure input

PredRBR [85], COACH [61], PRIME3D2D [86], RsiteDB [87, 88], and DeepSite [62] use structural information to report binding sites. The accuracy of NucleicNet can be improved (compared to protein structure only) if structural information of both interaction partners is provided.

Although PredRBR [85] works with PDB structures, users depend on the pre-trained model or need to train their own with a respective dataset of structures [85], which is why we excluded this tool from our evaluation. COACH combines multiple algorithms into one approach with a trained SVM classifier to determine consensus binding site residues of a given protein [61]. The tool also allows for a sequence-only input instead of a PDB structure, in which case a 3D model of the protein will be generated prior to the binding site residue prediction. The output lists and visualizes the results of the individual algorithms as well as the combined template-based COACH approach, with the top-ranked models and their calculated confidence score (ranging from 0 to 1) and binding residues [61]. PRIME3D2D refers to structure templates for prediction as well, using TMAlign [89] for protein and LocARNA [90] for RNA alignment to a template to build a RNA-protein complex model, followed by scoring of the potential binding site [86]. RSiteDB [87] focuses on extruded RNA nucleotides not involved in RNA base pairing, which could interact with protein binding pockets. It stores known nucleotide binding sites in RNA-protein structures, as well as provides a prediction service based on the data [88]. The algorithm predicts potential binding sites by determining atomic contacts between the PDB chains, dinucleotide patterns of the RNA, and (geometric) properties of a binding site [88]. DeepSite [62] is a knowledge-based deep convolutional neural network (DCNN) approach that predicts binding sites in proteins for different ligands and includes features such as atom types and chemical properties. The structures are treated like 3D images and ideally cover both the protein and ligands (RNA in our study). The potential residues are calculated and visualized in so-called ‘binding site centers’ which are supported by a score between 0 and 1.

Our literature search uniquely found PredPRBA [91] to predict the released energy after binding as a measure of possible interaction. The method uses multiple gradient boosted regression tree models for different classes of RNA-protein complexes in the dataset, with features extracted from both sequence and structure [91]. 3dRPC [92] is specifically focused on performing docking analyses of RNA and protein molecules into a scored complex. Docking is an alternative computational approach, which was expanded from protein-protein interactions to deal with RPIs only recently [93]. The challenge here lies mostly in the folding flexibility of RNA sequences and the implementation of a scoring function for RNA-protein interactions [93, 94].

PRince [95] reports the binding residues of both RNA and protein and produces an output in PDB format that can be used to visualize and explore the structural specificity of the RPI, focused on the accessible surface areas and the highlighted interface region. PRIHotscore [96] and OPRA [97] use the structural information from their respective PDB-derived datasets to report individual interaction probabilities per residue. PRIHotscore approaches the prediction via *in silico* alanine-scanning, which allows to identify interface amino acids as hotspots for RNA binding with assigned interaction scores [96]. OPRA on the other hand assigns interaction propensities based on the ratio of residue composition at RPI interfaces compared to that in the structures’ surface using statistical potentials [97].

PLIP [64] finds non-covalent interactions in binding sites between macromolecules or proteins and other biomolecules, like nucleic acids and ligands. Predictions are rule-based on the interaction geometry (i.e., distances and angles) between the residues in a given structure complex, and the physicochemical properties of the amino acids and nucleobases. PLIP reports a detailed list and visualizes the complex with probable interactions, their types, and involved residues and atoms.

#### Evaluation of *de novo* RPI prediction tools

Unfortunately, not all of the 30 described RPI prediction tools could be evaluated. As mentioned, GraphProt, iDeepS, and PredRBR still depend on pre-trained models or user’s HTS datasets, which is why they were excluded from the evaluation completely. We covered the remaining 27 algorithms, of which multiple ones provided no results to evaluate, due to different individual reasons: (i) not applicable to our examples (no matching data available): RBPmap; (ii) not installable/usable: aPRBind, IPMiner; (iii) computations did not finish or web server did not provide feedback about the status: GraphBind, PRIME3D2D, 3dRPC; (iv) no results are displayed or downloadable: PRince; (v) non-solvable error during input submission: beRBP, RsiteDB.

We introduce an evaluation score for our study, ranging from 0 (no evaluation criterion fulfilled) to 4 (all criteria fulfilled). Since the prediction tools vary widely in their amount and format of output, we decided on the criteria mentioned in the Methods to capture the tool’s usefulness for users as best as possible. The nine algorithms with no results to evaluate (listed above) received no assigned evaluation score. The full evaluation table is deposited at GitHub and includes input, parameters, runtime, output and further information for all tools for our analysis.

#### Evaluation dataset: Prediction results for four different biological RPIs

The interaction between the LARP7 protein and 7SK RNA is well-described [98] and functions for 7SK stability *in vivo* and assists in a stable association of the 7SK ribonucleoprotein (RNP) complex [106]. The xRRM domain at the C-terminus of LARP7 specifically binds the 3’-terminal U-rich stretch in the 7SK RNA and the top of stem-loop 4 (SL4) [98]. The base G312 and amino acids Y483 and R496 constitute the core interface (Fig. 2A). Additionally, the residues in the three structural elements *β*2 (RNP3 motif), *β*3, and *α*3 of LARP7 are important for the interaction. The main interaction types are hydrogen bonding and stacking interactions.

**Figure 2:**
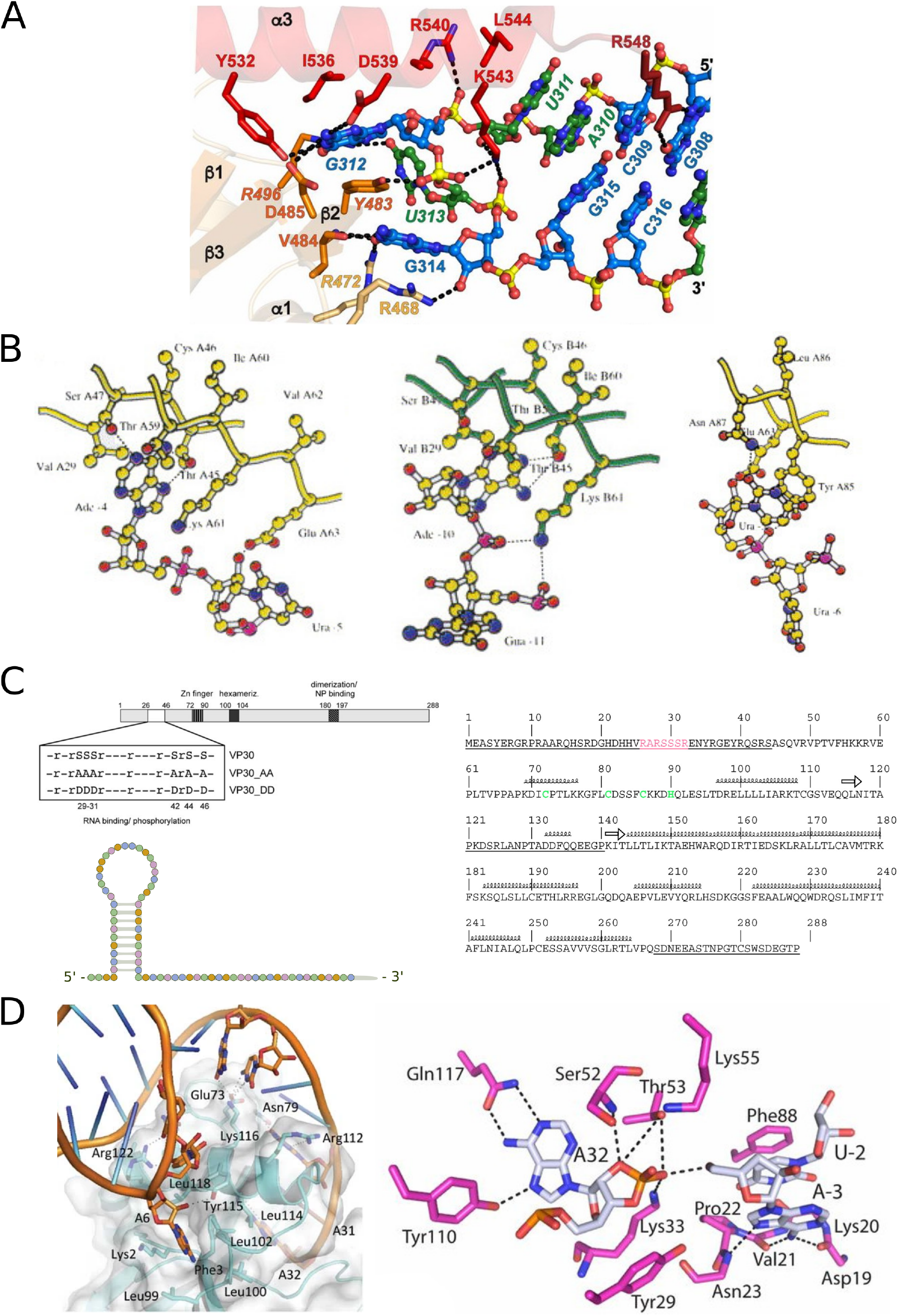
Known interactions for the four investigated RPI examples based on the respective literature. **(A)** The C-terminus of LARP7 (xRRM domain) binds the RNA with key amino acids in three structural elements (*β*2, *β*3, *α*3). The 7SK RNA is bound at the stem-loop 4 (SL4) and a U-rich region at the 3’-end. Adapted from [98]. **(B)** A 19-nucleotide-long RNA region in the MS2 phage genome folds into a hairpin structure bound at exposed positions by a dimer of the MS2 phage coat protein. The contacting amino acids are primarily conserved within the *β*-strands of the protein structure [99]. Adapted from [100]. **(C)** The N-terminal region of the Ebola virus VP30 protein binds the 3’-leader region of the RNA genome strand at nucleotides 80-54 with an arginine-rich region [101]. The optimal RNA substrate is single-stranded, of 40nt length and mixed base composition [102]. Further single amino acids contribute to the RPI [103]. Adapted from [101–103]. **(D)** The RPIs in the bacterial ToxIN system assist in assembling the heterohexameric complex of three toxI RNA pseudoknots and three ToxN monomers. The interactions are described to be focused around a few key nucleobases bound by various amino acids of the protein in multiple pockets. Adapted from [104, 105].

The relevant interacting regions for LARP7 and 7SK were predicted most detailed by PLIP and PRIHotscore (evaluation score of 4 and 3, respectively, Fig. 3). Both managed to positively predict the RPI involved residues while correctly excluding non-interacting ones. Since the available PDB complex represents the specific RPI region well, other tools considering structure also performed well with scores of 2: COACH, NucleicNet, and OPRA, as well as MEME using sequence only. The most important residues are predicted by multiple tools, but rarely the RPI is covered exhaustively or in full detail. For the protein residues, we observed many false positive predictions, i.e., amino acids predicted as interacting although not explicitly being mentioned as such in the corresponding literature [98]. This can be seen for example with aaRNA, hybridNAP, KYG, and catRAPID. For the latter, interactions are focused at the 5’-end of the 7SK RNA and across the complete LARP7 sequence, with an overall high interaction propensity of 84. As LARP7 binds to the UUU-3’OH and the SL4 stem-loop [107], the prediction at the 5’-end is not supported by the literature. Moreover, *in vivo*, the 5’-end of 7SK RNA is bound by the methylphosphate capping enzyme (MePCE) [98], denying the opportunity for interaction with LARP7 in this region. Another crucial aspect in prediction for structure-based tools is the presence of the RNA molecule in the 3D complex. Exemplarily, since the LARP7 PDB entry does not include the 7SK RNA structure, the results of DeepSite are less reliable than if the RNA was present, i.e., only few of the true positive amino acids for interaction are predicted. In the case of a model-based approach like COACH, the prediction is highly dependent on the availability of adequate models for the example at hand. The top-ranked model for LARP7 is based on a Polypyrimidine tract-binding protein PTB (PDB:2ADC, covers an RRM domain) with a CUCUCU-RNA strand as a ligand. Since the interacting stem-loop of 7SK consists of the motif AUGAUG, the U-richness might be reflected in the model but only allows limited conclusions to the interaction of LARP7 with 7SK RNA. PRIdictor only predicted a few interacting residues with its web application but not the web server, therefore getting a score of 0.5.

**Figure 3:**
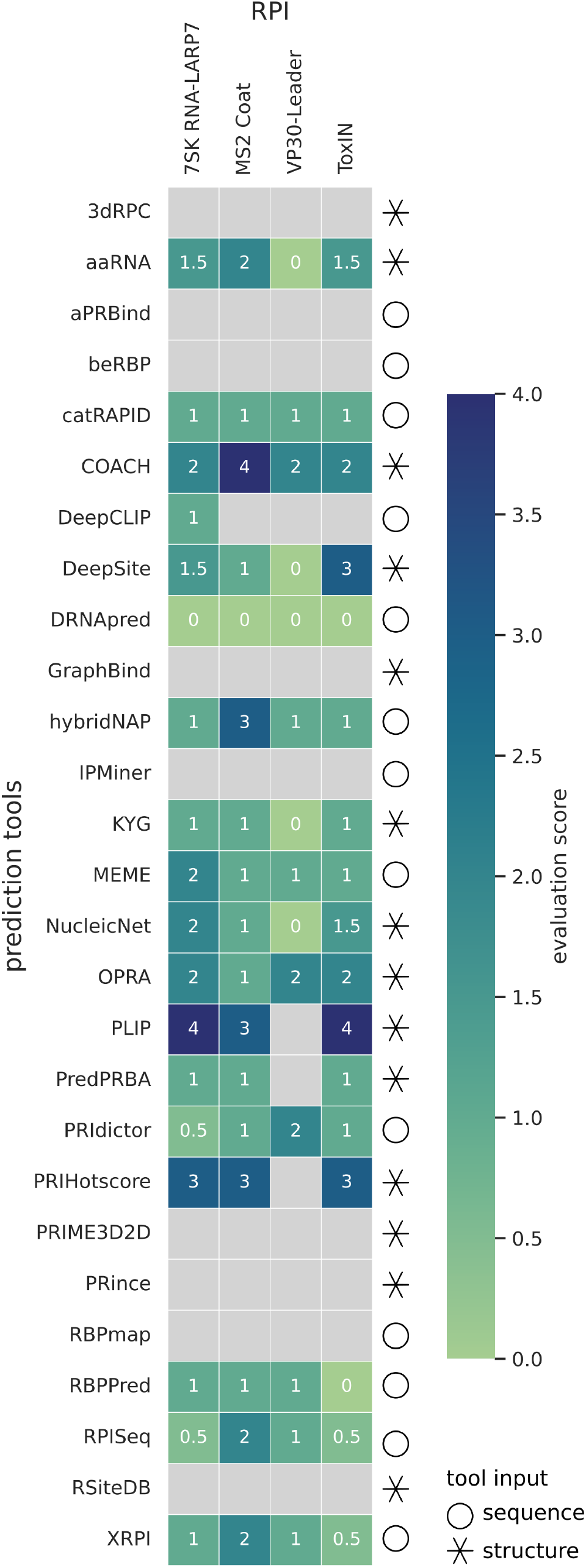
Evaluation heatmap. Based on the criteria listed in the Methods, we evaluated the RPI prediction tools for all four biological examples. The score therefore can range from 0 (lowest/worst) to 4 (highest/best) (color scale), or not assigned (gray) because of the reasons listed in section “Evaluation of *de novo* RPI prediction tools”. Tools marked with a circle require sequence input and the star indicates a structure input.

Another well-studied RPI example is the coat protein of the RNA phage MS2. As a dimer, it interacts with a 19-nucleotide-long RNA region in the phage genome, which folds into a hairpin structure and contains the initiation code for the replicase gene (Fig. 2B). The interaction of RNA and protein leads to a switch from replication to virion assembly in the viral life cycle [99, 108].

The interaction was detected most reliably and correctly by COACH, hybridNAP, PRIHotscore, and PLIP (Fig. 3). All of them correctly predict many of the RPI-involved amino acids [100] in the protein with high probability. However, they also predict some false positive residues, which cannot be distinguished without knowing the ’ground truth’ (hybridNAP, COACH), or miss a few important interaction partners (PRIHotscore, PLIP). With a score of 2, aaRNA, RPISeq, and XRPI perform slightly better for this compared to the other biological examples. aaRNA predicts binary binding propensities above threshold for some but not all interacting amino acids, and both RPISeq and XRPI confidently deliver high interaction probabilities with both classifiers (SVM: 0.96, RF: 1.0) or models (RPI2825: 0.9969, RPI390: 0.9191), respectively.

Our third RPI is the N-terminal region of VP30 in Ebola virus binding to a stem-loop structure in the 3’-leader region of the single-stranded, negative-oriented RNA genome at nucleotides 80-54 and the complementary antigenomic strand (Fig. 2C). The RPI is focused on an arginine-rich region in the protein and regulates the transcription of the virus [102, 103].

The best performing tools here, with a score of 2, were COACH, OPRA, and PRIdictor (Fig. 3). The latter does predict a subset of important interacting residues, but no regions as a whole, and also covers many false-positive hits. COACH and OPRA mostly do not provide RPI residues, which is in line with the input structure not including the main interacting region and was therefore the expected outcome. The algorithms of catRAPID, hybridNAP, MEME, RBPPred, RPISeq, and XRPI all predict some kind of interaction or motif based on the input sequence(s), however it does not represent the literature-known residues [102, 103] (score 1). Multiple structure-based tools cannot predict the RPI due to the disadvantageous PDB complex. This leads to the algorithms either reporting interactions despite the PDB entry not covering the interaction-important N-terminal region of the protein (score of 0: KYG, aaRNA, NucleicNet, DeepSite) or not being able to start their computations at all because of the lacking RNA molecule in the structure (no score: PLIP, PredPRBA, PRIHotscore).

As a fourth example, we investigated the bacterial toxin-antitoxin system ToxIN, which encodes an ABI (abortive infection) system in multiple, especially enteric bacteria [109]. To neutralize the toxin protein (ToxN), three ToxI RNA pseudoknots form a heterohexameric complex with three ToxN monomers mediated by RPIs in different binding pockets [104]. The interactions are focused around multiple amino acids, which group together to bind a few nucleobases, respectively (Fig. 2D).

Here, DeepSite, and again PRIHotscore and PLIP perform the best with scores of 3 and 4 (Fig. 3). PLIP correctly identifies nearly all interaction-involved residues for both RNA and protein and their respective relations. DeepSite and PRIHotscore perform similarly, but are missing some relevant amino acids. Both COACH and OPRA provide multiple false positive or false negative matches, respectively, in addition to some of the literature-known amino acids [104]. The two classifiers of RPISeq as well as the two models of XRPI come to different interaction probabilities each, which led us to assign half a point for both tools. RBPPred did predict the ToxN sequence as non-interacting, as opposed to its results for the three other examples.

## Discussion

### Quality (and quantity) of predictions differs greatly between different tools

The investigated RPI prediction tools work with different algorithms and on different data. Thus, their results vary vastly in their informative value and level of detail. Working with sequence input, the results of RBPPred, XRPI, and RPISeq provide a general single score or probability with no information on specific residues or regions. This might prove useful in case the user just needs this distinction of how likely an interaction is, e.g., for an “all-against-all” approach to compare multiple RNA and protein sequences. For RPISeq no obvious relation between the two classifiers is stated, complicating the interpretation whether an RPI can be trusted in the case of two diverging predicted probabilities. In such instances, the user cannot know if the RNA and protein are likely to interact, especially if there is no ground truth available. The motif finder MEME overall covers interacting regions in our evaluation, however the motifs are distributed across the sequences in general, challenging the motif’s specificity. Although MEME does not predict RPIs directly, it might prove useful if a motif of interest is known before analysis or to narrow down a potential amount of possible interaction sites for further experiments [69]. For probabilities or scores of individual residues in the sequence users can refer to DeepCLIP, hybridNAP, DRNApred, PRIdictor, or catRAPID. The latter additionally correlates every residue of both the RNA and protein sequence against each other. DRNApred does not predict any interactions for any of our examples and should therefore be used with caution.

Since spatial factors play crucial roles in RPIs [24, 25], structure-based tools generally produce more detailed results. The binding free energy calculated by PredPRBA provides an overall tendency for the RPI complex of interest. For per-residue-information, users can apply KYG, aaRNA, PRIHotscore, or OPRA, all of which supplement this with visualization in the structural complex. Specific binding sites are reported with NucleicNet, DeepSite, and COACH. The former highlights binding areas of potential interaction, whereas the latter two further indicate the respective amino acids involved. The multiple result models provided by COACH give the user the option to consider predictions with (potentially) low confidence scores, which may still prove useful for discovering “hidden” RPI-relevant amino acids. Without a ground truth, the manual evaluation of multiple models might be prone to errors. PLIP is the only one with information on both RNA and protein residues in its output, and also shows the types of interactions and therefore grants the most detailed predictions for all examples.

For all tools, data availability is essential. While sequence data is far more abundant than structural data for RPI complexes, the latter is mandatory for structure-dependent tools. A general problem in structure determination is the flexibility of the RPI complex [22], which is typically captured in one state only and thus might not reflect all aspects of the interaction. Working with datasets of bound and unbound states of the RBPs could support RPI prediction in this aspect. Additionally, only the protein structure of an RPI complex without the RNA partner is commonly determined, which impedes the structure-based algorithms to predict the RPI accurately. Furthermore, as seen in the Ebola virus VP30 example, similar issues arise when the protein structure is incomplete, especially when domains or regions important for the interactions are missing. This underlines the need for more structural data from experimental determination methods and continuous improvement of computational tools to predict as much accurate information as possible with as little data as necessary.

Here, we focused on *de novo* prediction tools instead of tools using or analyzing experimental RPI datasets (e.g. CLIP-based). The latter usually requires an additional training before the actual prediction to fit the model to the organism of interest. Especially for non-bioinformaticians, building and training a model on robust data sets is a challenging task and thus hinders the usage of such tools. Accessibility and usability are key factors in providing interpretable *in silico* RPI predictions.

### No tool is more accurate across all kingdoms

We examined the RPI prediction tools with RPI examples from different species: LARP7 and 7SK RNA acting in humans, MS2 phage coat protein and an RNA hairpin in the MS2 phage genome, VP30 protein and viral RNA leader region of Ebola virus, and the bacterial toxin-antitoxin system ToxIN. We did not observe drastic accuracy differences between the investigated tools in one of the examples. Some algorithms seemed to perform better for the MS2 phage coat protein RPI, namely hybridNAP, aaRNA, XRPI, and COACH, while others were challenged by the VP30 RPI, as seen for KYG, aaRNA, NucleicNet, PredPRBA, and DeepSite. Problems with the structure-based prediction for VP30 most likely stem from the PDB entries covering only parts of the protein and no RNA at all. Furthermore, the structure-based tools PLIP, PredPRBA, and PRIHotscore depend on the RNA structure in the PDB complex for their predictions. Including all structural components of the RPI generally grants more sound predictions, but is hindered if there is no adequate data available for the user to apply. Overall, in this investigation, prediction accuracies depended more on the functionality and input of tools (sequence versus structure) than the origin of biological examples. However, due to the restricted availability of training data, some tools, such as the proposed framework by Shulman-Peleg *et al*. [87], might be biased towards human or model organism RPIs or RPIs that are easier to investigate with the standard experimental methods. Consequently, these tools might perform more reliably for such RPIs, but might not generalize on non-model organisms and underrepresented RPIs. Even predictions within the organisms used for training models may fail to generalize for RPI instances, as with catRAPID (trained on human RPIs) and the human LARP7 and 7SK RNA example. RBPmap, RsiteDB, and beRBP are trained on human data as well, but since we could not evaluate results from those tools, we cannot deduce whether they would have been biased in favor of the human example. Similarly, algorithms providing pre-trained models for prediction are based on the available data and therefore focused on human data as well. For example, in our evaluation DeepCLIP only performs for the 7SK RNA, since LARP7 is the only protein available as a pretrained model. Furthermore, the restricted data availability extends also to the input for the tools, as mentioned above. The higher quality and amount of information the input data has, the more accurate the prediction tools can calculate potential RPIs, regardless of the species or kingdom.

### Post-processing potential of RPI prediction tools

Besides the prediction results themselves, the tools differ regarding how well their output can be used for further downstream analyses. Only displaying the results at the web server without the possibility for download of important output data hinders the user from meaningfully integrating a prediction tool into a self-build pipeline. We ranked NucleicNet, PRIdictor, catRAPID, and DeepSite as tools with a low post-processing potential. The annotated PyMol [110] session file provided by NucleicNet and the heatmap plot by catRAPID are downloadable, but only represent the predictions visually without the potential to parse the data. Besides the plot visualizations, PRIdictor only provides a text file for the protein predictions, not the RNA partner, which does not represent the result data in a well-retrievable way. DeepSite lets the user download the spatial coordinates of the ‘interaction centers’, not the interacting residues themselves.

Many tools provide their RPI prediction results in one or multiple parsable file formats, in addition to visual representations, depending on the respective tool. This includes DeepCLIP, MEME, hybridNAP, DRNAPred, aaRNA, COACH, KYG, PLIP, PRIHotscore, and OPRA. RBPPred, RPISeq, XRPI, and PredPRBA result in just a score value, which is downloadable as a text file for the latter two.

We cannot make statements regarding the post-processing potential for RBPmap, aPRBind, GraphBind, IPMiner, beRBP, PRIME3D2D, RsiteDB, PRince, and 3dRPC, as these tools either did not complete their calculations, did not process the input, or it was not possible to get them running (see Results).

## Conclusion

RPIs are ubiquitous in all life forms and can be studied with experimental detection methods and bioinformatic prediction algorithms based on their interaction features. With the growing amount of available data, current models and approaches can be improved on or expanded with this data to support RPI research and understanding further. In recent years, many tools and workflows have been proposed to predict RPIs with and without the requirement of HTS data. While this is a positive development for the field, users might lose the overview of accessible tools fitting their needs.

In this study, we provide an overview of RPI prediction tools and proposed a guide tree. We structured this overview and tree according to the tools’ required input and the degree of detail produced by their output. With this evaluation, we provide a guide for users to support the identification of appropriate tools for their research.

To assess the reliability of tools utilizing machine- and deep-learning techniques, we investigated the algorithms in more detail using four known RPI examples covering different kingdoms of life. The tools report varying amounts of detail and information about the specifics of the respective interaction. By having a ground truth at hand, this study gives insights into the reliability and interpretability of the tools. Computational predictions always have to be evaluated carefully, and this study is not exhaustive in terms of possible RPI examples and the performance of tested tools. Without prior knowledge, not all tools are valuable for *de novo* prediction use cases. Low confidence of the algorithms complicates interpretation of results or separation of false positives and false negatives from true predictions. The structure-based tools overall provided more details on the interactions, but rely on the availability of RNA-protein structure complexes.

Due to limited RPI data, predictions might be biased toward the interaction mechanisms of the organisms used for training and benchmarking. Many available tools rely on training data originating from human or model organism data. To overcome these issues, homology-based approaches, including evolutionary information, as intended in COACH or aaRNA, may be suited. However, additional and continuous refinement of such models is needed when new data is available. Thus, to deepen our understanding of RPIs and their exact mechanisms, we expect to see a continuing close exchange between experimental detection and *in silico* prediction models in the future, i.e., experimental data being fed into existing or new algorithms, while *in silico* prediction results reduce the space of potential RPI interactions to investigate in the lab and thus the necessary time and costs.

## Materials and methods

### RPI data selection

We used four cross-species examples for the RPI tool evaluation: (i) human protein LARP7 with the 7SK snRNA, (ii) MS2 phage coat protein with an RNA hairpin in the phage’s genome, Ebola virus VP30 protein with the viral RNA leader region, and (iv) bacterial toxin-antitoxin system ToxIN. The examples are based on how well they are researched, whether there is available data to use for the tools, and to cover species from different kingdoms.

For algorithms requiring a 3D structural complex, we provided a corresponding PDB entry structure for each example (see Table 1). For the 7SK RNA-LARP7-complex, we chose the PDB:6D12 structure entry because it specifically contains the interacting regions of RNA and protein. However, DeepSite could not resolve the necessary features of this structure (error for protonation of amino acids), which is why we used PDB:5KNW (same as PDB:6D12 but without RNA) for this tool instead. Both structures only represent the relevant RNA-interacting xRRM domain of LARP7 and not the whole protein structure. Determined structures for VP30, unfortunately, currently only cover the protein partially (5DVW starts with position 139 compared to the UniProt sequence), and lack RNA molecules. The preferred PDB entry for the ToxIN complex was 2XD0 which comprises the full heterohexameric molecule, but for KYG and NucleicNet, we had to choose PDB:4ATO (only one protein and RNA monomer each) due to complications of these tools with the former structure.

**Table 1:**
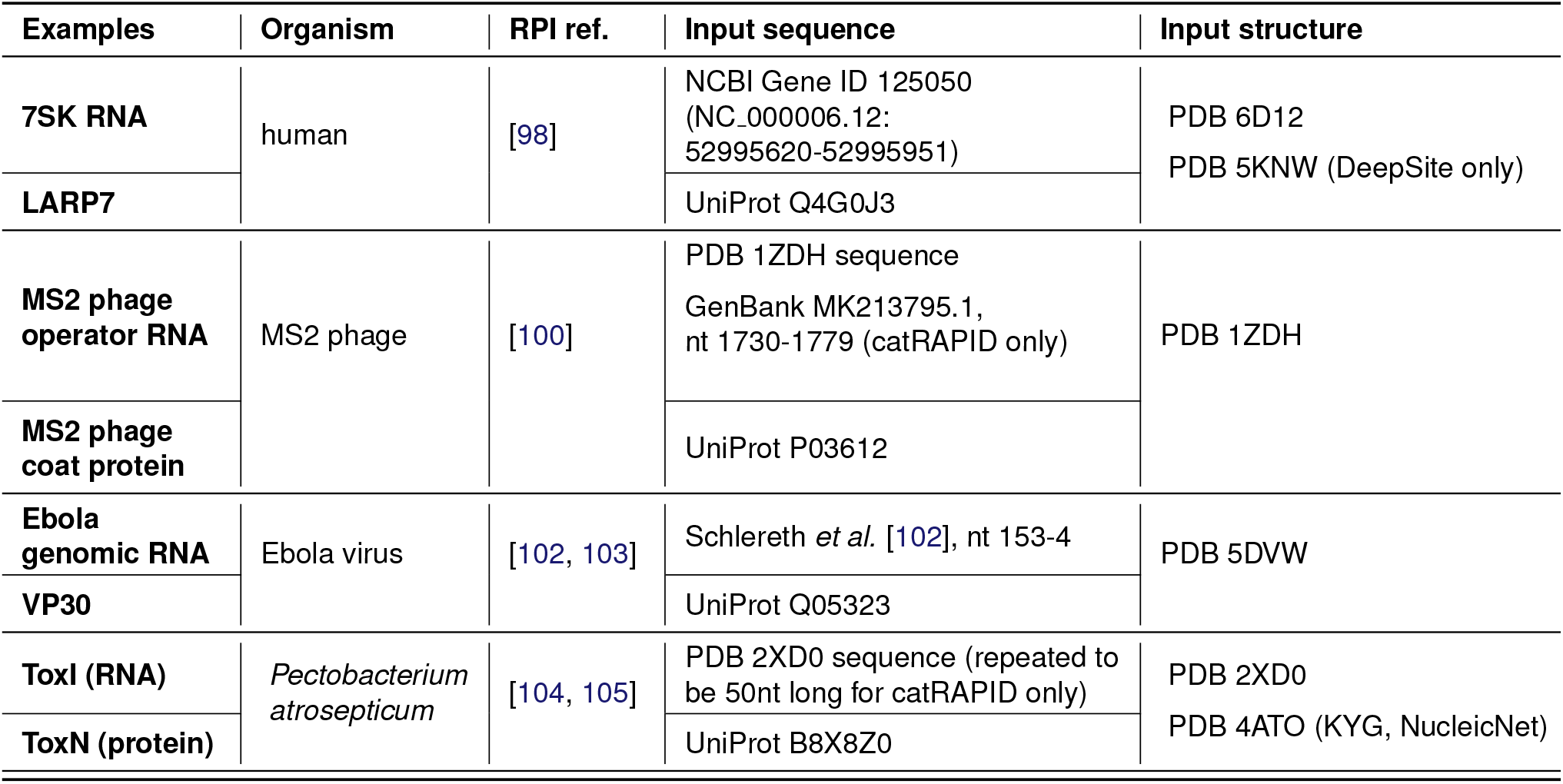
Overview of RPI examples and respective input for the tools. The table lists the RPI examples used in this study and the literature describing known interacting residues. Furthermore, it shows which data (sequence/structure) was used for the prediction tool evaluation.

For the sequence-based tools, we downloaded the protein sequences from the respective UniProt entries (see Table 1). The sequence of the 7SK RNA corresponds to NCBI Gene ID:125050. For the MS2 operator RNA, we used the sequence from PDB:1ZDH (the wild-type U/T at position 11 has been substituted with C for improved interaction and crystallization process [100]). Since catRAPID needs at least 50nt for input, we used the corresponding GenBank entry (MK213795.1) to add 16 upstream and 15 downstream nucleotides to the sequence. We retrieved the interacting genomic, negative-oriented RNA strand of Ebola virus from the supplementary of Schlereth *et al*., covering nucleotides 153-4 [102], which contains the most important positions for the RPI. We picked the RNA sequence of ToxI from entry PDB:2XD0. We extended this entry for catRAPID by appending the first 14 nucleotides of the sequence to the 3’-end to reach 50nt total. We justify this with the biological function: The RNA sequence is present as multiple repeats in the bacterium before being cut into the interacting monomers [109].

### Evaluation of tool results

The details regarding web server or stand alone usage, as well as parameters, run-time, input, output, are listed in Table `tool_evaluation.xlsx` (GitHub repository). We evaluated the results with the support of information about the respective RPIs described in the literature, gained in experiments. The knowledge originates from introducing point substitutions in connection with the determination of the equilibrium dissociation constant (*K*_*d*_) using ITC (isothermal titration calorimetry) [98], crystallization experiments and structure determination [98, 100, 104, 105], deletion mutants and site-directed mutations [102, 103], or EMSA analysis [102].

We decided on the following evaluation criteria:

- Did the algorithm predict any interaction for the given input?
- Does the prediction cover the correct region (in protein and/or RNA)?
- Does the prediction cover the correct interacting residues?
- Does the tool report a trustable confidence score (*>*=70%, if applicable)?

Because of the great differences of format and amount of output provided by the algorithms, selection of these criteria proved difficult. Fulfillment of each criterion leads to 4 points total maximum. A score of zero implies that the tool did provide prediction results, but none of the evaluation criteria were fulfilled. No score implies that the tool was not applicable to the respective example, did not complete its computations, or could not be installed. Table `tool_evaluation.xlsx` (GitHub repository) gives an overview of the tool evaluation and relevant information. The evaluation heatmap was plotted with an in-house python script and finalized with InkScape v.1.1.2 [68].

Furthermore, we assessed the potential for post-processing the results of each tool, e.g., for downstream analysis after the RPI prediction. (+) denotes parsable, downloadable results the user can potentially incorporate into a computational pipeline. Tools with a (-) provide viewable content on their web server, but no option for download or potential to incorporate the results directly into downstream analyses of the user. (n/a) marks the tools which were not applicable (no evaluation score) and we thus could not assess those regarding their post-processing potential.

## Supporting information

Supporting information can be found at GitHub.

## Acknowledgements

We thank Dr. Emanuel Barth and Prof. Dr. Manja Marz for proof-reading the manuscript.

